# T cell exhaustion dynamics in systemic autoimmune disease

**DOI:** 10.1101/2023.12.23.573167

**Authors:** Christi N. Turner, Juan Camilo Sanchez Arcila, Noah Huerta, Avi Rae Quiguoe, Kirk D.C. Jensen, Katrina K. Hoyer

## Abstract

Unlike in infection and cancer, T cell exhaustion in autoimmune disease has not been clearly defined. Here we set out to understand inhibitory protein (PD-1, Tim3, CTLA4, Lag3) expression in CXCR5- and CXCR5+ CD8 and CD4 T cells in systemic lupus erythematosus. CXCR5+ CD8 and CD4 T cells express PD-1 and engage B cells in germinal center reactions, leading to autoantibody formation in autoimmunity. We hypothesized that CXCR5+ CD8 T cells develop an exhausted phenotype as SLE autoimmunity expands from initial to chronic, self-perpetuating disease due to chronic self-antigen exposure. Our results indicate that there is no exhaustion frequency differences between sexes, although disease kinetics vary by sex. CXCR5+ CD8 T cells express primarily IFNγ, known to promote autoimmune disease development, whereas CXCR5-CD8 T cells express TNFα and IFNγ as disease progresses from 2-6 months. Tim3 is the highest expressed inhibitory marker for all CD4 and CD8 T cell populations demonstrating potential for terminally exhausted populations. CTLA4 expression on CD4 T cells suggests potential tolerance induction in these cells. We identified exhaustion phenotypes within autoimmune disease that progress with increasing lupus erythematosus severity and possibly provide a feedback mechanism for immunological tolerance.

**Highlights:**

1. CXCR5- and CXCR5+ CD8 T cells expand with rate of disease in SLE mouse model.
2. CXCR5+ CD8 T cells are low contributors to TNFα disease progression unlike CXCR5-CD8 T cells but may increase disease mechanisms through high IFNγ production.
3. Inhibitory markers upregulate in frequency with the highest amounts seen in Tim3+ populations. Tim3+Lag3+ expression may be an indicator of terminal differentiation for all populations.
4. Inhibitory marker expression frequency was unrelated to sex.

## Introduction

Autoimmunity affects millions of individuals throughout the world and is induced through the inappropriate recognition of self-antigen by the immune system. While many studies have focused on the mechanisms behind autoimmune disease development, activation and T cell driven disease outcomes, much remains unknown. Within chronic antigen environments, such as cancer, viral infection and autoimmune disease, T cells can become functionally “exhausted” losing their proliferative capacity and cytotoxic effector production, memory formation and possibly interactions with B cells. The role that T cell exhaustion plays in autoimmunity is particularly unclear as only a handful of studies have explored exhaustion in rheumatoid arthritis, type I diabetes, psoriatic lesions, alopecia areata and systemic lupus^1–7^.

T cell exhaustion characteristics have been largely defined through models of infection and cancer^8–10^. Exhausted T cells can vary in their inhibitory progression from progenitor to terminally exhausted^8^. Terminal exhaustion (Tex) in T cells is determined by loss of IL-2 signaling and subsequent downregulation of TNFα, reduced proliferation, loss of effector molecules such as perforin and granzyme b, and upregulation of IL-10 and multiple inhibitory receptors such as programmed cell death protein 1 (PD-1) and Tim3^9,11,12^. PD-1, originally discovered as an indicator of thymic activation, is an early inhibitory protein along with other proteins of inhibition, such as Tim3, Ctla4 and Lag3^13,14^.

We previously discovered CXCR5+ CD8 T cells within IL-2 knockout mice and other systemic antibody-mediated autoimmune diseases^15^. These T cells are capable of interacting with B cells, promoting antibody-class switching and plasma cell differentiation in systemic autoimmunity^16,17^. CXCR5+ T cell populations that express characteristics of pre-exhausted (Tpex) and terminally exhausted (Tex) subsets arise during lymphocytic choriomeningitis virus (LCMV), hepatitis B virus (HBV), simian immunodeficiency virus (SIV) and friend virus infections^18–24^. CXCR5- and CXCR5+ CD8 and CD4 T cells upregulate exhaustion genes during rapid, systemic autoimmune disease in IL-2 deficient mice^15^. This exhaustion gene signature identified by RNA sequencing led us to explore CXCR5+ CD8 T cell exhaustion kinetics in a slower progressing autoimmune disease mouse model of systemic lupus erythematosus (MRL/lpr).

Systemic lupus erythematosus (SLE) is an autoimmune disease characterized by expanded immune cell populations and sex-based differences in disease severity^25^. We hypothesized that CXCR5+ CD8 T cells develop an exhausted phenotype as SLE autoimmunity expands from initial to chronic, self-perpetuating disease. We observed CXCR5+ CD8 T cell expansion in MRL/lpr mice over the course of autoimmune disease. As CXCR5+ CD8 T cells expand, multiple inhibitory receptors are upregulated at varying times with increased TNFα, IFNγ, and granzyme b but decreased perforin with disease progression. However, CXCR5+ CD8 T cells express less exhaustion receptors than CXCR5-CD8 and CXCR5-CD4 counterparts. Overall, CXCR5+ CD8 and CXCR5-CD8 T cells express increasing levels of exhaustion receptors as disease progresses from 2-6 months. This progressive inhibition may be a feedback mechanism to subdue autoreactivity in chronic autoimmune disease. Alternatively, exhaustion receptors may upregulate on healthy bystander, non-self-reactive T cells as a mechanism to maintain a non-activated state.

## Materials and Methods

### Animals

MRL/MpJ and MRL/lpr mice were purchased from Jackson Laboratories and donated as a gift from Dr. Gabriela Loots at Lawrence Livermore National Laboratory. BALB/c IL-2 knockout were used with littermate IL-2 wildtype or heterozygous (WT) controls (^26^, GSE112540). All animal protocols and procedures were followed according to IACUC standards and the UC Merced Department of Animal Research Services.

### Lymphocyte collection and flow cytometry

Lymph nodes and spleens were dissected and disassociated through wire mesh then processed in phosphate buffered saline supplemented with 1% fetal bovine serum (FBS; Omega) then red blood cells lysed in ammonium chloride lysis buffer for 1 min at room temperature followed by washing. Lymphocytes and splenocytes were counted with a hemacytometer and antibody-labeled for flow cytometry. All antibodies were purchased from eBioscience unless noted otherwise. Cells were first labeled with CXCR5-biotin (SPRCL5) for 1 hr at RT, then CD4-PerCPCy5.5 (RM4-5), CD8-APC-eFluor780 (53-6.7), PD-1-Fitc (J43), tim3-APC (8B.2C12), lag3-PE (C9B7W; Biolegend), fixable viability 506, CD11b-PECy7 (M1/70), CD11c-PeCy7 (N418), ly6G-PeCy7 (RB6-8C5), B220-PeCy7 (RA3-6B2), CD49b-PeCy7 (DX5) for 25 min at 4°C. Cells were fixed with foxP3 fixation/permeabilization kit (Thermofisher/Invitrogen) following manufacturer guidelines for intracellular staining of CTLA4-PE610 (UC10-4B9) and streptavidin-BUV395 (SA; BD Biosciences). Data was acquired by flow cytometry on a Becton Dickinson LSR-II and analyzed using FCS Express with Diva version 7 (DeNovo Software). Lymphocytes were gated based on the forward scatter area (FSC-A) and side scatter area (SSC-A). Doublets were removed and cells were gated on singlets and viable cells (viability 506). Viable cells were gated on CD4+ or CD8+ then CXCR5+ or CXCR5-.

### Cell Stimulations

2×10^6^ cells were plated per well in a 24-well plate in RPMI media supplemented with penicillin, streptomycin, glutamate, non-essential amino acids and sodium pyruvate. Cells were stimulated with 0.2 ng/mL phorbol 12-myristate 13-acetate (PMA), 2 ng/mL ionomycin, and secretion blocked with 10 μg/mL brefeldin-A (Invitrogen) for 5-6 hours at 37°C with 5% CO_2_. Cells were collected after incubation and antibody-labeled for flow cytometry. Cells were first stained for CXCR5-biotin (SPRCL5) for 1 hr at RT, then stained for CD4-PerCPCy5.5 (RM4-5), CD4-e450 (GK1.5), CD8-APC-eFluor780 (53-6.7), B220-PECy7 (RA3-6B2), CD11b-PECy7 (M1/70), CD11c-PECy7 (N418), ly6G-PECy7 (RB6-8C5), CD49b-PeCy7 (DX5), fixable viability 506 for 25 min at 4°C. Some cells were tertiary stained for IL-21R-Fc (149204; R&D Systems) for 25 min at 4°C. All cells were fixed with foxP3 fixation/permeabilization kit (Thermofisher/Invitrogen) for intracellular staining of IL-10-PerCPCy5.5 (JES5-16E3), IL-21-PE (FFA21), IFNγ-APC (XMG1.2), IL-1B (NJTEN3), IL-2-PE (JES6-5H4; Biolegend), TNFα-Fitc (MP6-XT22), perforin-Fitc (eBioOMAK-D), granzyme b-PE (NGZB), and streptavidin-BUV395 (SA; BD Biosciences). Flow cytometry was completed on a Becton Dickinson LSR-II. Flow cytometry analysis as performed using FCS Express with Diva version 7 (DeNovo Software).

### CD107a Degranulation Assay

Cells were stimulated with PMA and ionomycin according to the above parameters in the presence of CD107a-PerCP710 (eBio1D4B) at 37°C with 5% CO_2_ for 5 hours; brefeldin-A was added after 1 hour of incubation. Flow cytometry was completed on a Becton Dickinson LSR-II and data analyzed using FCS Express with Diva version 7 (DeNovo Software).

### PCA

Principal component analysis (PCA) was calculated to evaluate cell population distribution dependent on age, sex and genotype using a log10 transformed matrix with the function princomp from the package stats^27^. First, we used a PCA to filter variables with a potential contribution to explain the spatial distribution of the data. We chose cell populations with loading scores higher than the average (>1.14) of the loading values in Dim1 and these variables were used to perform a clustering analysis. To find the variables with the highest contribution to explain the spatial distribution of the data on Dim1 and Dim2, we used the function fviz_contrib(). The cell populations selected for having the highest contribution were used to plot the bidimensional distribution of the individuals by age, sex and phenotype using the function fviz_pca_ind from the package factoextra aiming to understand the association of age, sex, and genotype. In the PCA plots, the length of each arrow denotes the contribution of each variable to explain data distribution and the angle between arrows for each pair of variables determines the correlation between variables in the bidimensional space (0° is equal to complete correlation, 90° = no correlation and 180° = negative correlation). Only Dim1 and Dim2 data was considered in the analysis. To confirm the observations in the PCA plots, we evaluated the contribution of each variable to explain the data distribution in the multivariate space by examining the loadings along each principal component.

### Heatmap

Data was preprocessed in FCS Express with Diva version 7 (DeNovo Software) by gating on CD4 or CD8 T cells followed by CXCR5- or CXCR5+. Gated cell population frequencies were exported to .csv tables for posterior analysis in R. To evaluate the diversity of exhaustion markers on CD4 and CD8 subsets, we performed a PCA ordination and selected variables with Dimension1 loading scores > 1.14. The variables selected were used to study the clustering patterns associated with a particular genotype, sex, or age. A 2-dimensional heatmap was built using R package ComplexHeatmap^28^. The function hclust from the package fastcluster^29^ using a log10 transformed distance matric build from the data was used for running the hierarchical clusters. Clustering methods Euclidean distance and ward.D2 were used.

### Statistics

GraphPad Prism (Version 8) was used to compute statistics. Student’s t-test was performed to assess differences between two means. ANOVA with Tukey correction was used to compute differences between multiple means where appropriate. Statistical measurements are indicated in each figure legend and described as NS = not significant, *p < 0.05, **p < 0.01, ***p ≤ 0.001, or ****p ≤ 0.0001.

## Results

### CXCR5- and CXCR5+ CD8 T cells expand with SLE rate of disease

We began our investigation by exploring the changing landscape of CXCR5-versus CXCR5+ CD8 T cells across 2-6 months of age in MRL/lpr mice relative to MRL/MpJ controls. Peak of systemic lupus erythematosus (SLE) in MRL/lpr mice is 4 months of age with end life occurring at approximately 5-6 months for females and 6-8 months for males^25,30,31^. This age range covers T cell responses from early (2 months), mid (4 months) and late (6 months) disease time points. CD8 T cell frequency in male lymph nodes is decreased compared to controls across 4-6 months, but increased in total cell number, due to lymphoproliferation in these mice (Figure 1A). A similar trend in CD8 T cell frequency and total number is observed in lymph nodes of female mice (Figure 1B), with similar numbers and frequency between males and females of each genotype. As expected, CXCR5+CD8 T cell population is smaller than CXCR5-CD8 T cells in male and female mice^18^. CXCR5- and CXCR5+ CD8 T cell numbers begin to expand by 4 months in males and 2 months in females in association with disease progression (Figure 1C-F). Similar cell frequencies in males and females were identified in spleens across 2-6 months with smaller total CXCR5- and CXCR5+ CD8 T cell populations (Suppl. Figure 2).

**Figure 1.**
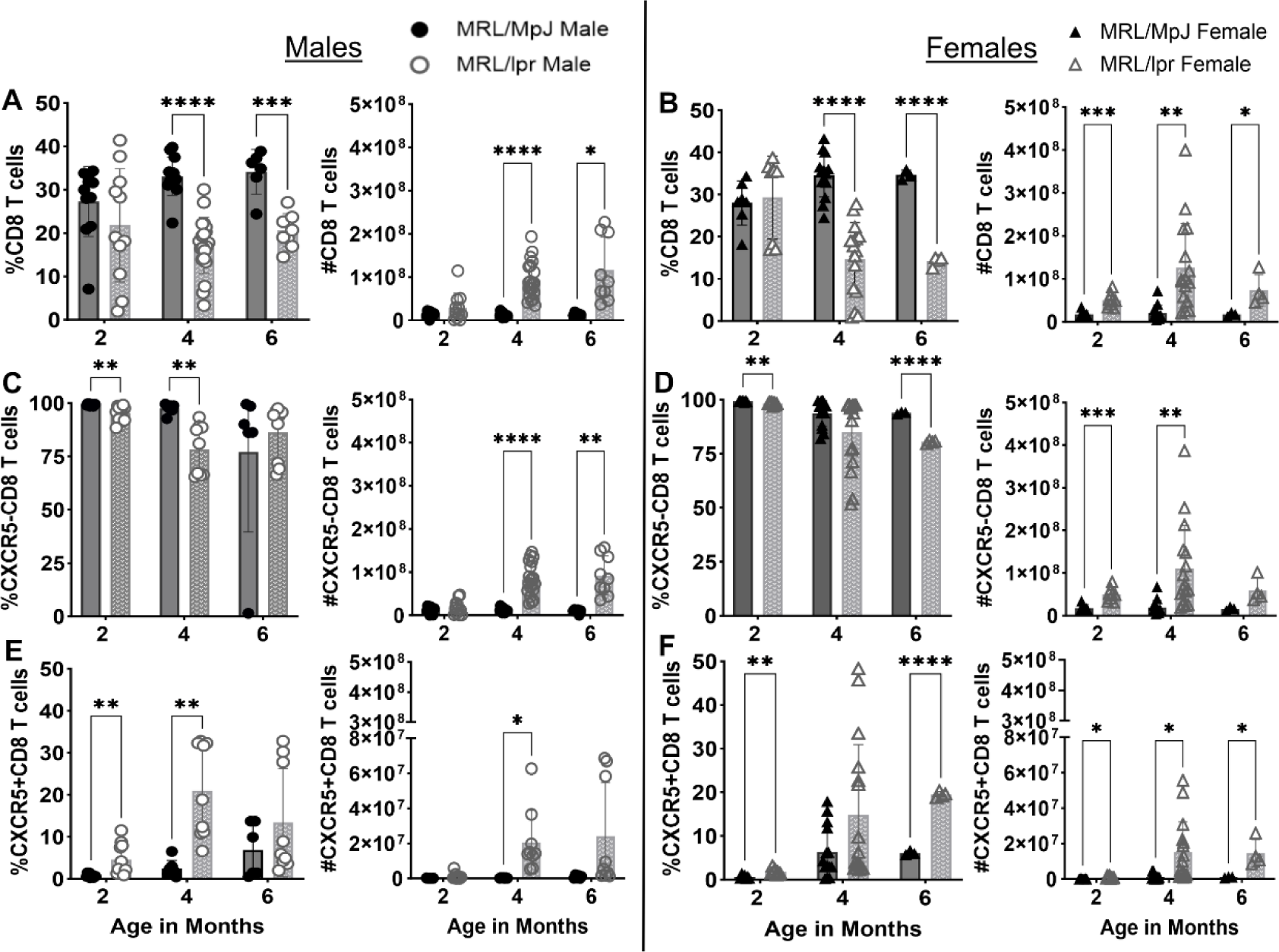
CXCR5- and CXCR5+ CD8 T cell expansion in SLE lymph node. (**A-B**) Frequency and total numbers of all CD8+ T cells across 2, 4, 6 months in male and female MRL/MpJ and MRL/lpr mice. (**C-D**) Frequency and total numbers of CXCR5-CD8 T cells across 2, 4, 6 months in males and female mice. (**E-F**) Frequency and total numbers of CXCR5+CD8 T cells across 2, 4, 6 months in males and female mice. All frequencies and total numbers determined by flow cytometry and cell counting. Each symbol represents one animal and data is representative of 1-5 independent experiments per comparison. Males are denoted by circles and females by triangles. Statistics were performed relative to indicated controls with unpaired Student *t* test. *p < 0.05, **p < 0.01, ***p ≤ 0.001, ****p ≤ 0.0001.

### CXCR5+ CD8 T cells produce IFNγ associated with stimulating germinal center development in SLE

MRL/lpr mice are known for variation in cytokine and effector molecule production that contribute to disease burden. Pro-inflammatory cytokine tumor necrosis factor alpha (TNFα) and interleukin-1 beta (IL-1β) promote disease severity in SLE patients and mice through multiple processes including germinal center organization and B cell antibody promotion^32,33^. Since CXCR5+ CD8 T cells are known regulators of germinal center reactions we next identified their cytokine capacity in the context of SLE following *ex vivo* stimulation. TNFα starts low and increases significantly in 6-month male CXCR5-CD8 lymph node T cells; in females TNFα remains slightly elevated across 2-4 months (Figure 2A and 2C; Suppl. Fig. 3). CXCR5+ CD8 T cells are not significant contributors of TNFα compared to CXCR5-CD8 T cells (Figure 2A and 2C; Suppl. Figure 3). IFNγ-producing CD8 cells significantly expand over the course of SLE in males and females (Figure 2A and 2C; Suppl. Figure 3). IFNγ production in CXCR5+ CD8 T cells rise slower and with lower frequency reaching maximum at disease endpoint (Figure 2A and 2C; Suppl. Figure 3). CXCR5-CD8 T cells have a greater change in IFNγ expression frequency across all time points in male and females and both peripheral organs than CXCR5+ CD8 T cells. A small percent of CXCR5-CD8 splenic T cells produce IL-1β while CXCR5+ CD8 T cells produce minimal IL-1β in the lymph node or spleen of male mice (Figure 2B and 2D; Suppl. Figure 3). No differences in IL-10 and IL-21 were observed across timepoints or organs though IL-2 increased at 4 months in male spleens (Figure 2B and 2D; Suppl. Figure 3B and 3D).

**Figure 2.**
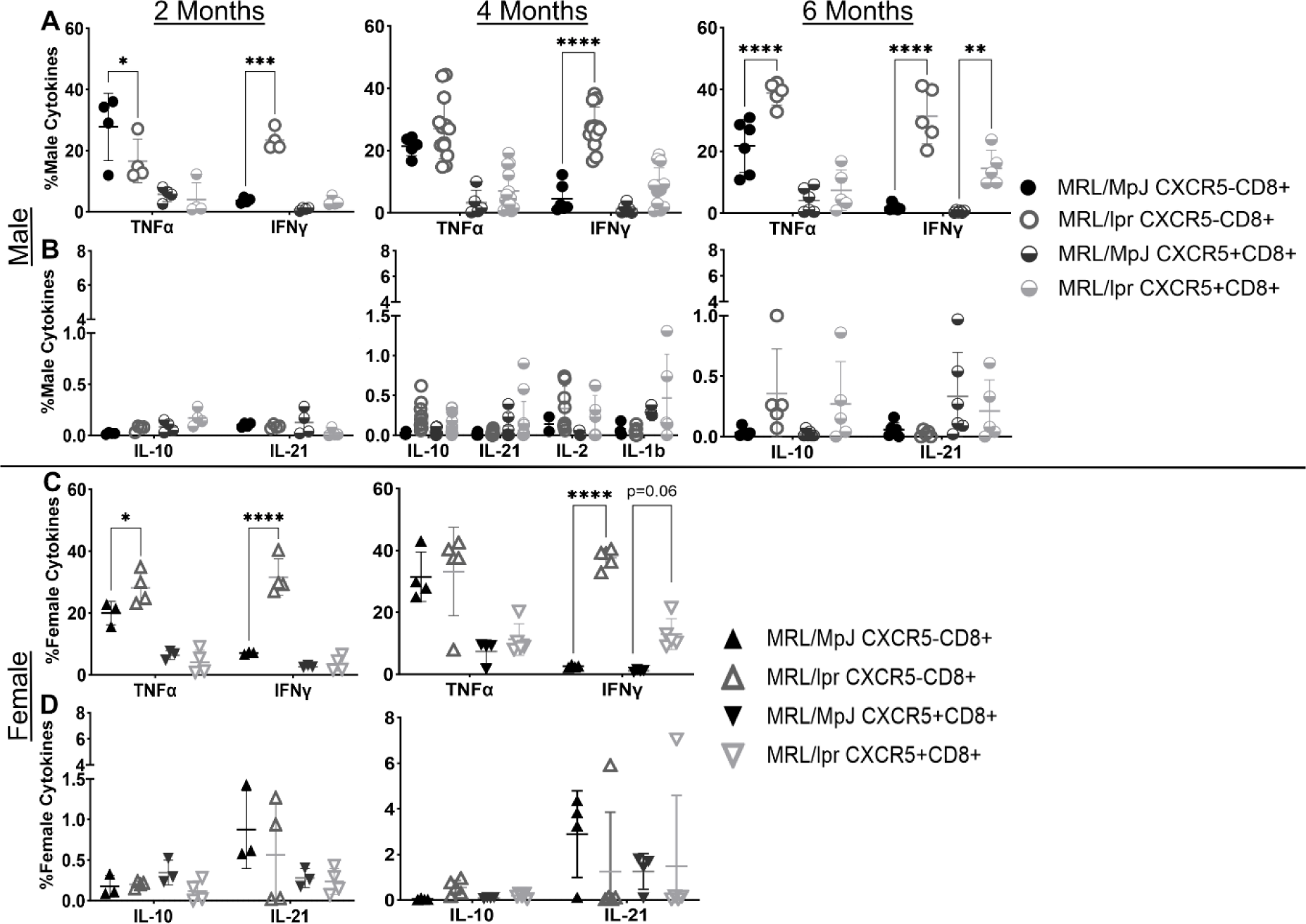
CXCR5- and CXCR5+ CD8 T cells influence SLE progression through different cytokines in lymph node. (**A**) Frequency of TNFα and IFNγ producing CXCR5- and CXCR5+ CD8 T cells from male MRL/MpJ and MRL/lpr mice at 2, 4, and 6 months of age. (**B**) Frequency of IL-10 and IL-21 producing CXCR5- and CXCR5+ CD8 T cells from male mice at 2-6 months of age. Frequency of IL-2 and IL-1β producing CXCR5- and CXCR5+ CD8 T cells from male mice at 4 months of age. (**C**) Frequency of TNFα and IFNγ producing CXCR5- and CXCR5+ CD8 T cells from female MRL/MpJ and MRL/lpr mice at 2, 4, and 6 months of age. (**D**) Frequency of IL-10 and IL-21 producing CXCR5- and CXCR5+ CD8 T cells from female mice at 4 months of age. All frequencies determined by flow cytometry. Each symbol represents a different animal and data is representative of 2-5 independent experiments per comparison. Males are denoted by circles and females by triangles. Statistics: ANOVA relative to indicated controls with multiple comparisons and Tukey correction. *p < 0.05, **p < 0.01, ***p ≤ 0.001, ****p ≤ 0.0001.

Cytotoxic effector molecules are decreased in SLE patients^34^, and perforin-deficient lupus mice develop accelerated autoimmune disease with increased autoantibody production^35^. We next investigated cytotoxic effector capacity of CXCR5- and CXCR5+ CD8 T cells in MRL/lpr mice. Perforin production is almost non-existent in both CD8 T cell populations in male and female lymphoid organs at all time points (Figure 3A-C, E-G; Suppl. Figure 4A-C, E-G). Granzyme B frequency and total numbers significantly increase in females at 4 months in both peripheral organs but decrease in male lymph nodes (Figure 3A-C, E-G). Splenic CXCR5-CD8 T cells significantly upregulate granzyme B from 2-4 months with increased total CXCR5+ CD8 T cells expressing granzyme B at 6 months (Suppl. Figure 4A-C, E-G). We next assessed CD8+ T cell degranulation via CD107a following stimulation with phorbol 12-myristate 13-acetate and ionomycin. CXCR5-CD8 T cells showed slight increases in CD107a frequency and total numbers from 4-6 months in males and 2-4 months in female lymph nodes (Figure 3D and 3H). CD107a also increased with disease kinetics in females from 2 to 4 months (Suppl. Fig 4H). CXCR5+ CD8 T cells were not powerful producers of CD107a in SLE.

**Figure 3.**
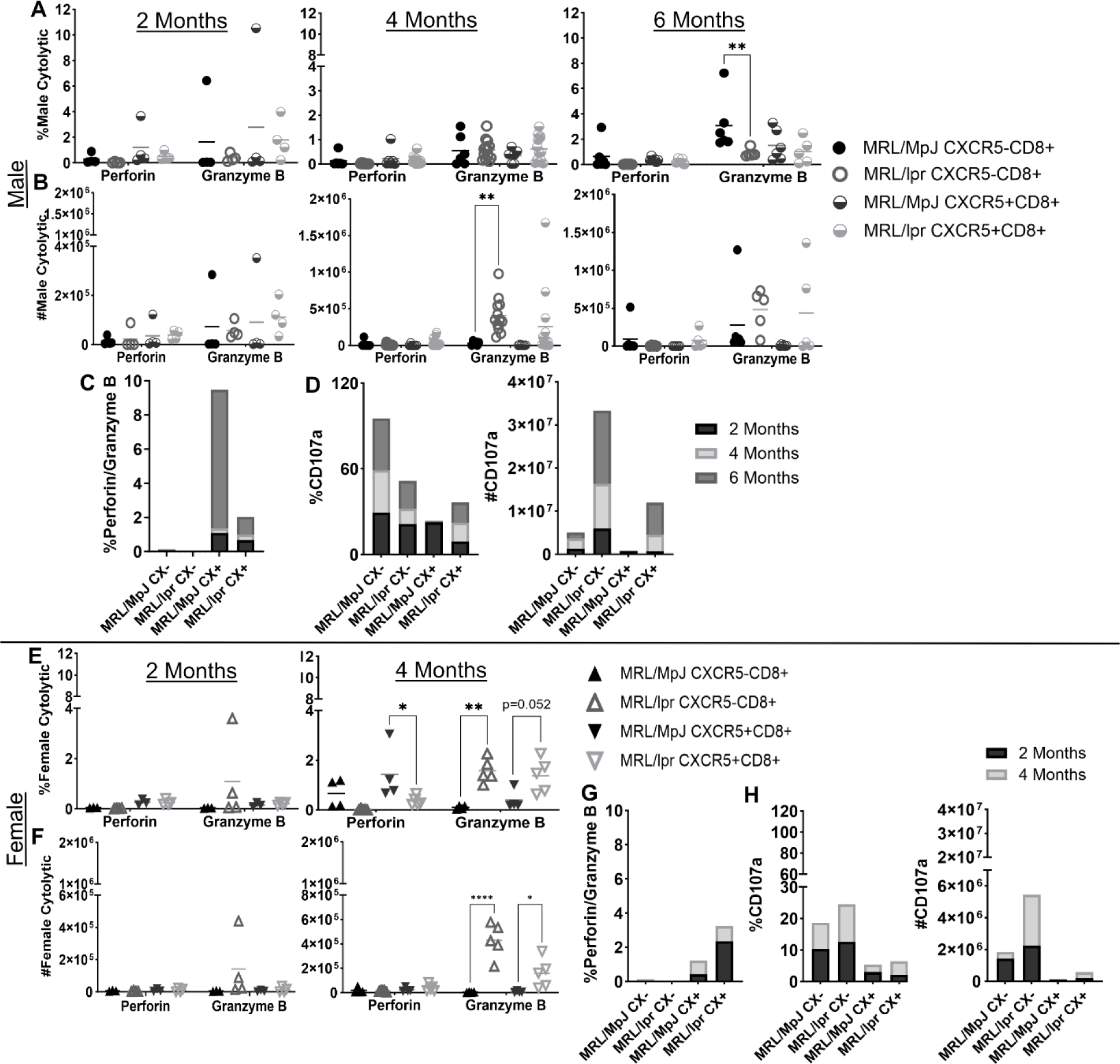
Female CXCR5- and CXCR5+ CD8 T cells produce higher granzyme B, but not perforin. (**A-B**) Frequency and total numbers of perforin and granzyme B producing CXCR5- and CXCR5+ CD8 T cells from male MRL/MpJ and MRL/lpr mice at 2, 4, and 6 months of age. (**C**) Amount of perforin to granzyme B production in male MRL/MpJ versus MRL/lpr CXCR5- and CXCR5+ CD8 T cells. (**D**) Frequency and total number of CD107a degranulation in male MRL/MpJ and MRL/lpr CXCR5- and CXCR5+ CD8 T cells. (**E-F**) Frequency and total numbers of perforin and granzyme B producing CXCR5- and CXCR5+ CD8 T cells from female MRL/MpJ and MRL/lpr mice at 2, 4, and 6 months of age. (**G**) Amount of perforin to granzyme B production in female MRL/MpJ versus MRL/lpr CXCR5- and CXCR5+ CD8 T cells. (**H**) Frequency and total number of CD107a degranulation in female MRL/MpJ and MRL/lpr CXCR5- and CXCR5+ CD8 T cells. Frequency and total numbers determined by flow cytometry and cell counting. Each symbol represents a different animal and data is representative of 2-5 independent experiments per comparison. Males are denoted by circles and females by triangles. Statistics: ANOVA relative to indicated controls with multiple comparisons and Tukey correction. *p < 0.05, **p < 0.01, ****p ≤ 0.0001.

### Exhaustion markers reveal declining functional capacity in CXCR5- and CXCR5+ CD8 and CD4 T cell populations

CD8 T cells upregulate inhibitor proteins in infection and cancer resulting in effector exhaustion. A stepwise co-expression of multiple inhibitory receptors characterizes progression of T cell exhaustion. To explore inhibitory marker expressions in CXCR5- and CXCR5+ CD8 and CD4 T cells during autoimmune disease we assessed expressions of PD-1, Tim3, CTLA4 and Lag3. PD-1 expression in CXCR5- and CXCR5+ CD8 T cells is variable between mice but increases from 2-4 months in MRL/lpr females while remaining consistent in MRL/lpr males from 2-6 months (Figure 4A-C; Suppl. Figure 5A-B). CXCR5-CD8 T cells expressing Tim3 increase across all time points, sex, and peripheral organs (Figure 4D-E; Suppl. Figure 5C). CXCR5+ CD8 T cells reveal increased Tim3 frequency that is most notable in the female spleen at 2-4 months (Figure 4F; Suppl. Figure 5D). Elevated Tim3 in CXCR5-CD8 T cell frequency suggests that these cells may be terminally exhausted during autoimmunity in MRL/lpr mice (Figure 4E).

**Figure 4.**
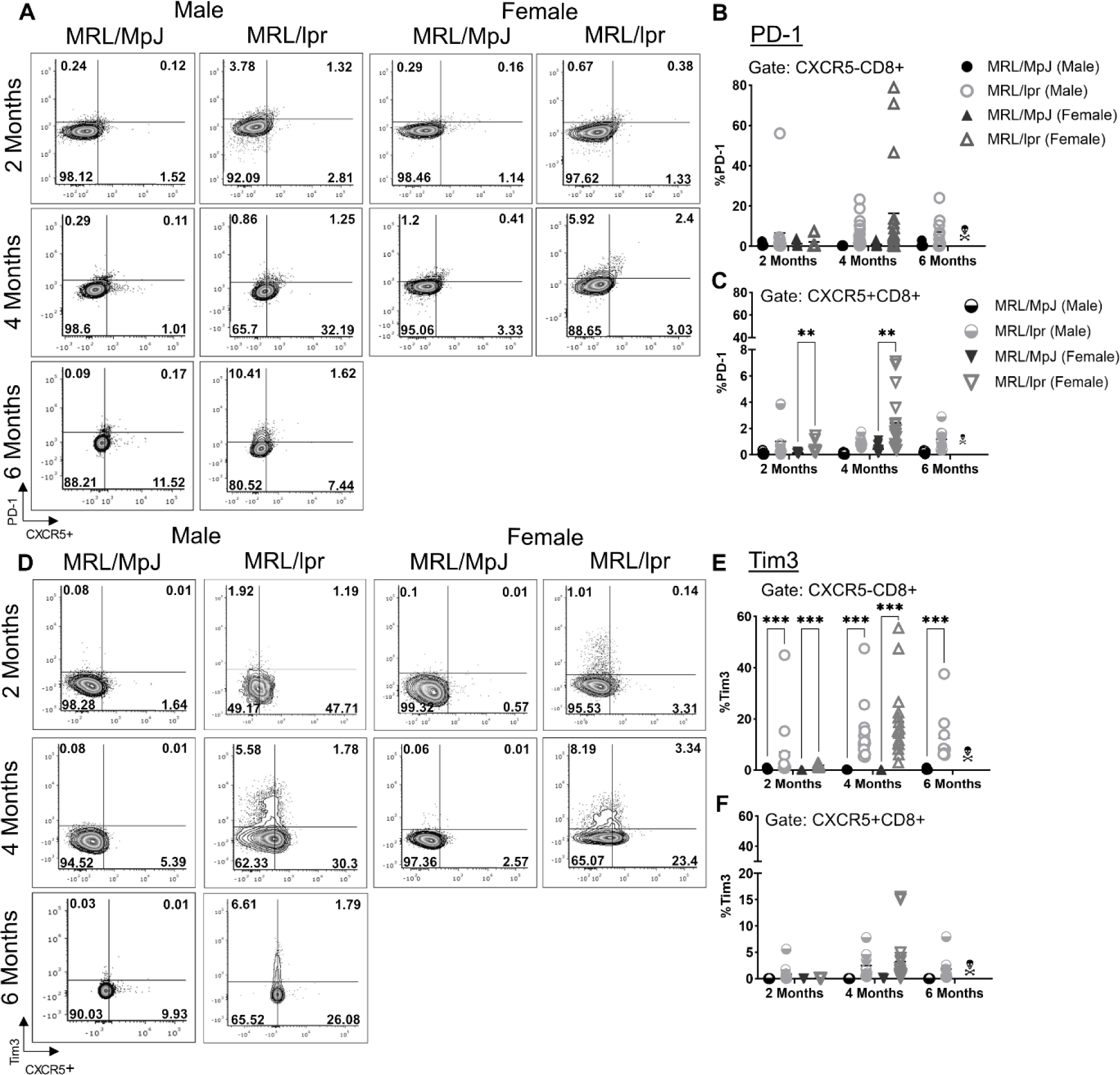
PD-1 and Tim3 expression on CXCR5- and CXCR5+ CD8 T cells from lymph node. (**A**) Representative flow plots comparing CXCR5 and PD-1 expression on live B220^−^CD11c^−^ CD11b^−^GR-1^−^ CD8 T cells from 2, 4, and 6-month male and female MRL/MpJ and MRL/lpr mice. (**B**) Frequency of PD-1 expressing CXCR5-CD8 T cells from 2, 4, and 6-month male and female MRL/MpJ and MRL/lpr mice. (**C**) Frequency of PD-1 expressing CXCR5+ CD8 T cells from 2, 4, and 6-month male and female MRL/MpJ and MRL/lpr mice. (**D**) Representative flow plots comparing CXCR5 and Tim3 expression on B220^−^CD11c^−^CD11b^−^GR-1^−^ CD8 T cells from 2, 4, and 6-month male and female MRL/MpJ and MRL/lpr mice. (**E**) Frequency of Tim3 expressing CXCR5-CD8 T cells from 2, 4, and 6-month male and female MRL/MpJ and MRL/lpr mice. (**F**) Frequency of Tim3 expressing CXCR5+ CD8 T cells from 2, 4, and 6-month male and female MRL/MpJ and MRL/lpr mice. Each symbol represents a different animal and data is representative of 2-5 independent experiments per comparison. Males are denoted by circles and females by triangles. Statistics: ANOVA relative to indicated controls with multiple comparisons and Tukey correction. **p < 0.01 and ***p ≤ 0.001.

Female, but not male, MRL/lpr CXCR5+ CD8 T cells also demonstrate increased frequency of CTLA4 expression at 2 and 4 months (Figure 5A, C). CTLA4 expressing CXCR5- and CXCR5+ CD8 T cells were identified across time points, sex, and peripheral organs in varying frequency (Figure 5A-C; Suppl. Figure 6A-B). Lag3 expression is upregulated in male lymph node CXCR5-CD8 T cells across all time points of disease (Figure 5D-E; Suppl. Figure 6C). CXCR5+ CD8 T cells expressed Lag3 across all time points and sex in lymph nodes and in the female splenic cells (Figure 5F; Suppl. Figure 6D). Next, we pooled all CXCR5- or CXCR5+ CD8 T cells expressing any exhaustion markers, whether single or overlapping, from males and females from each genotype across each time point. These graphs demonstrate that inhibitory marker expression exists at a greater frequency in MRL/lpr SLE mice compared to control MRL/MpJ mice (Figure 5G-H; Suppl. Figure 6E), and sometimes at high frequencies and expression level mean fluorescence intensity (data not shown).

**Figure 5.**
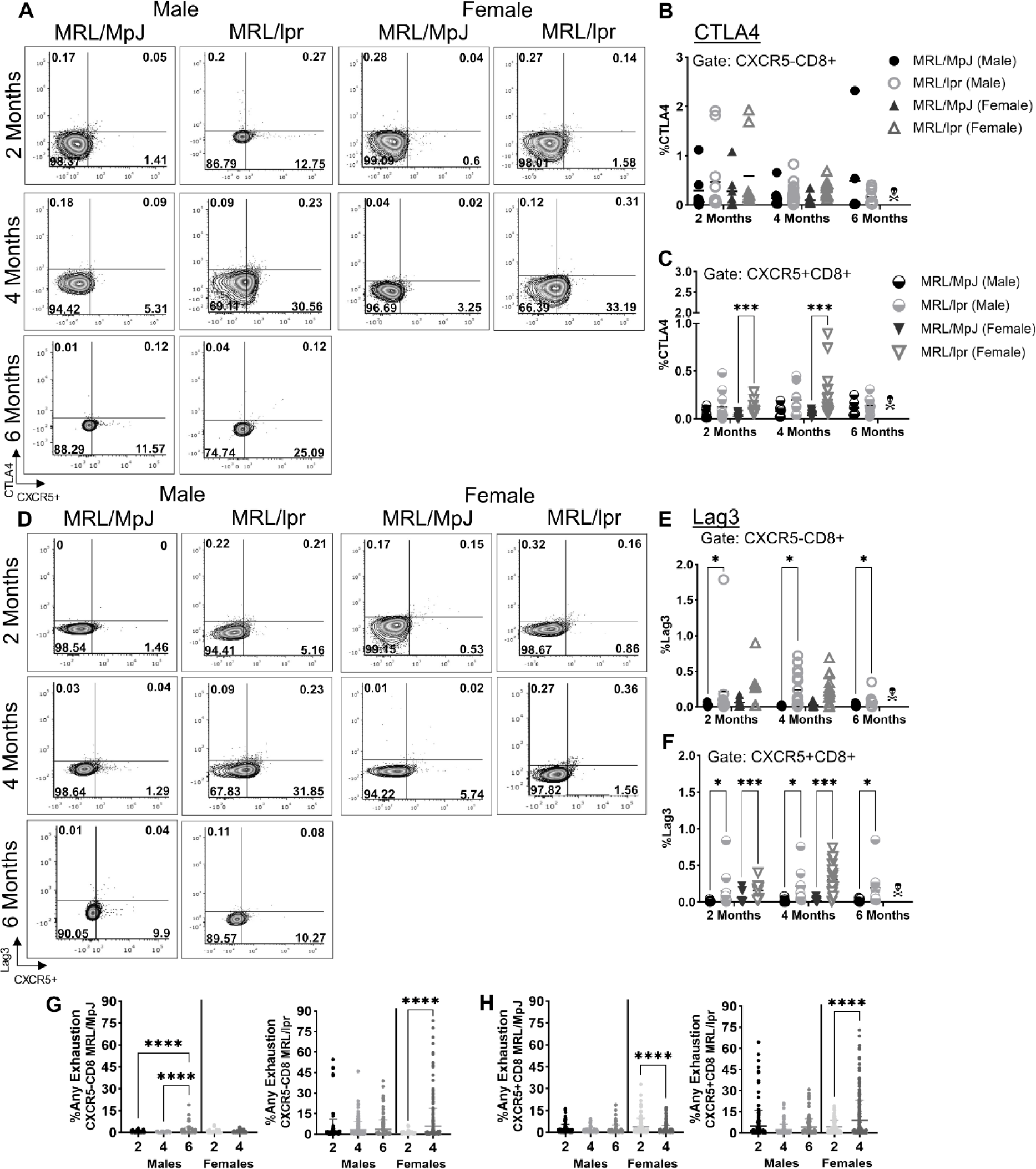
CTLA4 and Lag3 expression on CXCR5- and CXCR5+ CD8 T cells from lymph node. (**A**) Representative flow plots comparing CXCR5 and CTLA4 expression on live B220^−^ CD11c^−^CD11b^−^GR-1^−^ CD8 T cells from 2, 4, and 6-month male and female MRL/MpJ and MRL/lpr mice. (**B**) Frequency of CTLA4 expressing CXCR5-CD8 T cells from 2, 4, and 6-month male and female MRL/MpJ and MRL/lpr mice. (**C**) Frequency of CTLA4 expressing CXCR5+ CD8 T cells from 2, 4, and 6-month male and female MRL/MpJ and MRL/lpr mice. (**D**) Representative flow plots comparing CXCR5 and Lag3 expression on B220^−^CD11c^−^CD11b^−^GR-1^−^ CD8 T cells from 2, 4, and 6-month male and female MRL/MpJ and MRL/lpr mice. (**E**) Frequency of Lag3 expressing CXCR5-CD8 T cells from 2, 4, and 6-month male and female MRL/MpJ and MRL/lpr mice. (**F**) Frequency of Lag3 expressing CXCR5+ CD8 T cells from 2, 4, and 6-month male and female MRL/MpJ and MRL/lpr mice. (**G-H**) Any individual and double exhaustion marker frequencies were added together for each MRL/MpJ and MRL/lpr male and female mouse. The average frequency was graphed to show all exhaustion marker expression across 2-6 months in male and female mice. Each symbol represents a different animal and data is representative of 2-5 independent experiments per comparison. Males are denoted by circles and females by triangles. Statistics: ANOVA relative to indicated controls with multiple comparisons and Tukey correction. *p < 0.05, ***p ≤ 0.001, ****p ≤ 0.0001.

CD4 T cells are known to express inhibitory proteins though the regulation and purpose still remains unclear compared to CD8 T cells^9^. CXCR5+ CD8 T cells resemble CXCR5+ CD4 T follicular cells in surface marker expression and B cell interactions while retaining cytolytic potential^36–38^. Since CXCR5+ CD8 T cells can acquire CD4 Tfh functionality, we wondered if CXCR5+ CD4 T cells can behave like CD8 T cells regarding inhibitory receptor expression. We sought to explore and compare CXCR5- and CXCR5+ CD4 T cell exhaustion marker expression. Frequency of cells expressing PD-1 was elevated in female lymph nodes at 2 and 4 months in CXCR5-total numbers and frequency and total numbers of CXCR5+ CD4, but only in total numbers of CXCR5-CD4 T cells in spleen (Suppl. Figure 7A-D). Tim3 expression frequency in CXCR5-CD4 T cells expands across all time points, sex, and peripheral organs (Suppl. Figure 8A, C). CXCR5+ CD4 T cells show increased expression though not highly upregulated in lymph nodes, but in female spleens at 2 and 4 months (Suppl. Figure 8B, D). CTLA4 expression frequency expands in male and female lymph nodes and spleens across 2-4 months (Suppl. Figure 9A-D). CXCR5-CD4 T cell Lag3 frequency increases in males across 2-4 months in the lymph node, while female spleens upregulate Lag3 in CXCR5- and CXCR5+ CD4 T cells (Suppl. Figure 10A-D). The amount of upregulated inhibitory markers on both CXCR5- and CXCR5+ CD4 T cell populations is an indication that these cells are potentially losing functional capacity and becoming more dysfunctional as disease progresses from 2-4 months in females and 2-6 months in males (Suppl. Figure 11).

### Low amounts of terminal exhaustion may exist but is superseded by singular exhaustion marker expression in murine SLE

Progression of T cell exhaustion is defined by upregulation of multiple inhibitory markers including PD-1, Tim3, CTLA4, and Lag3. We performed a cluster analysis with a CXCR5- and CXCR5+ CD8 and CD4 T cells to explore the diversity of exhaustion markers in these subsets (Figure 6A). A clear clustering pattern was detected on the vertical data distribution associated almost exclusively with the phenotype of the studied individuals. MRL/MpJ mice were characterized by presenting intermediate frequency values of the exhaustion cell populations. In contrast, MRL/lpr presented high frequencies of CXCR5- and CXCR5+ CD8 and CD4 lymphocytes expressing exhaustion markers. A specific example of the contrasting distribution pattern can be illustrated with some cell populations located on the first cluster of the heatmap that presented high values on MRL/lpr and low frequencies such as MRL/MpJ of CXCR5+CD8+Lag3+Tim3+, CXCR5+CD8+CTLA4+Tim3+, CXCR5-CD4+Lag3+Tim3+, CXCR5-CD8+CTLA4+Tim3+, CXCR5-CD8+PD1+Lag3+, and CXCR5-CD8+Lag3+Tim3+. Also, it is important to notice that Tim3 was the most prevalent marker of exhaustion expressed by the cell populations included in the heatmap with 13 out of 19 cell populations.

**Figure 6.**
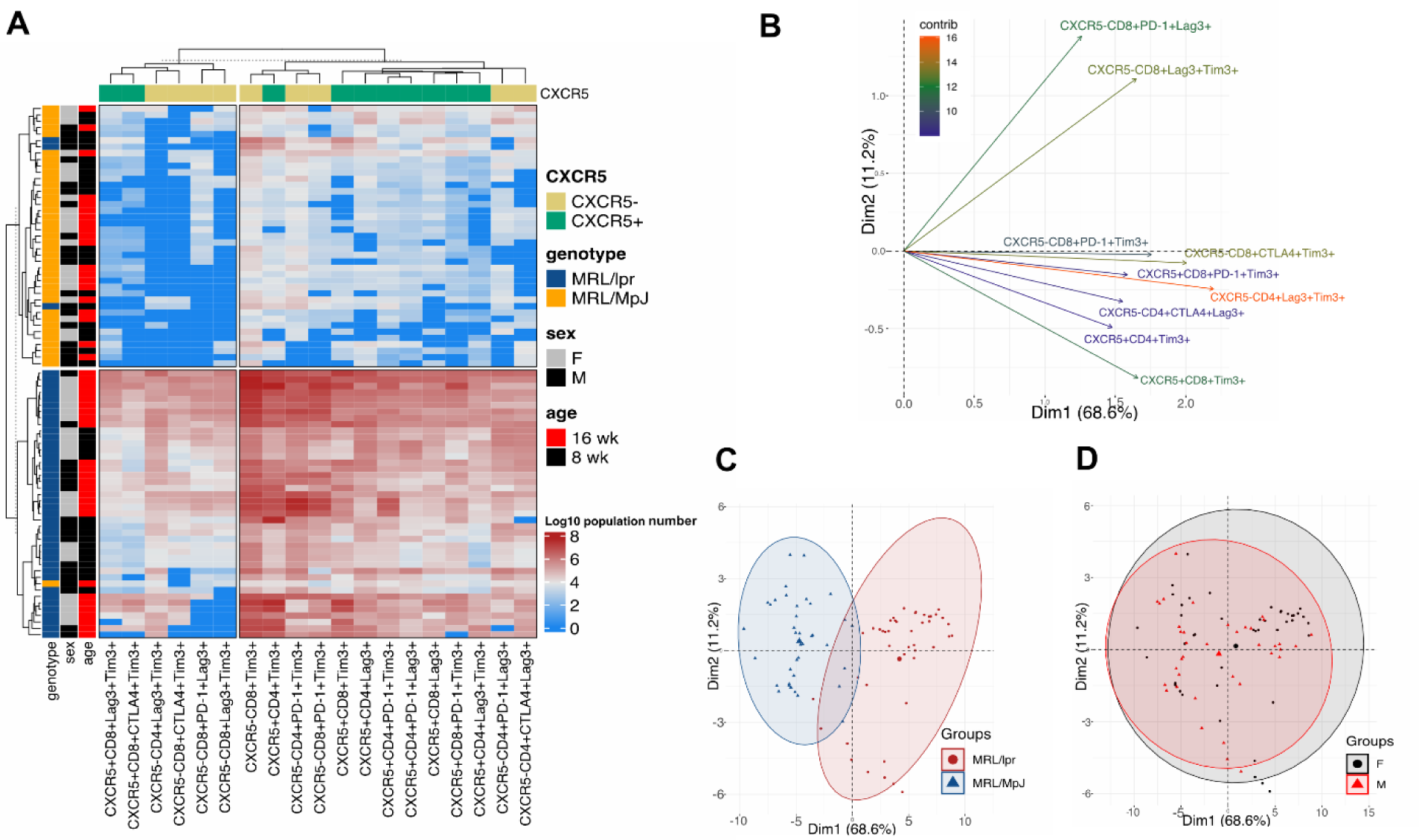
Multivariate analysis of T cell lymph nodes shows disease severity and not sex is the highest contributing factor to AD exhaustion. **(A)** Bidimensional heatmap of lymph node cell population Log10 numbers expressing exhaustion markers in CD4 and CD8 T cells. Each cell represents a unique Log10 value of a specific cell population for one individual. Vertical cluster represents grouping pattern for all the individuals from different genotypes (Dark blue = MRL/lpr, Dark yellow = MRL/MpJ), sex (gray = F, black = males), and age (black = 8 wk, red = 16 wk). Grouped horizontal clustering of cell populations with similar Log10 counts and CXCR5 expression (pale yellow = CXCR5-, green = CXCR5+). **(B)** PCA ordination of selected variables with a high contribution to the spatial distribution of individuals in bidimensional space. The Dim1 and Dim2 values represent the total contribution of selected variables to the spatial grouping for each corresponding dimension. The length of the arrow and heat color (highest = red, lowest = blue) are proportional to the degree of contribution of each variable to PCA 53 grouping pattern. PCA biplots with overlay ellipses showing individuals’ distribution pattern by genotype **(C)** or sex **(D).**

Because of the high diversity of cell populations, we decided to plot a PCA using data from the populations with the highest contribution for the spatial distribution of the data (Figure 6B). In total, Dim1 explained 68.6% of the total distribution of the data and Dim2 explained 11.2%, indicating that variables with the highest loading values along the Dim1 are responsible for most of the explanation of the data distribution. We noticed that nine cell populations were specially implicated with the pattern of data distribution, with a specific high contribution of CXCR5-CD4+Lag3+Tim3+ and CXCR5-CD8+CTLA4+Tim3+ cell populations. Also, we wanted to confirm the distribution pattern dependent on the mouse strain previously observed in the clusters/heatmap but using a PCA. We observed again that the data distribution was dependent exclusively on the genotype indicating differences in disease severity (Figure 6C) and not sex (Figure 6D).

## Discussion

This study set out to determine exhaustion marker upregulation in T cell populations of systemic autoimmune disease. We focused on comparing differences between CXCR5- and CXCR5+ T cells in MRL/lpr mice during disease progression based on cell expansion, cytokine and effector molecule production, inhibitory receptor upregulation and sex. CXCR5+ CD8 T cells in viral infections and cancer are associated with clearance of virally infected cells and tumor abatement, but in autoimmune disease they associate with disease severity through B cell interactions and antibody formation^16,19,20^. CXCR5- and CXCR5+ CD8 T cells expand with disease severity in systemic lupus erythematosus, a novel finding for CXCR5+ CD8 T cells in SLE^25^. SLE severity progresses through TNFα and IFNγ cytokine upregulation^39^. CXCR5-CD8 T cells expressed these inflammatory cytokines whereas few CXCR5+ CD8 T cells express TNFα, but express IFNγ at increasing rates as disease progressed from 2-6 months in males and females. Interestingly, the opposite has been observed in human tonsils and follicular lymphoma where CXCR5+ CD8 T cells produce high levels of TNFα demonstrating possible differences in diseases^37^. In some studies, type I interferons, such as IFNα, are the predominate cytokine expressed and initiate tissue damage by inducing type II interferon, IFNγ, from T cells^40–42^. Altogether this suggests that CXCR5+ CD8 T cells are minimally associated with the TNFα inflammatory response that initiated disease progression, but contributes more to disease burden and possibly autoantibody formation through IFNγ upregulation^41^. In contrast, CXCR5-CD8 T cells may promote disease initiation via TNFα production.

Follicular helpers are considered activated effectors that enter germinal centers and engage activation of B cells in peripheral organs^43–45^. SLE decreases effector T cell production of perforin and granzymes and CD107a has altered functionality in SLE patients that associates with inflammatory response instead of autoimmune remission or infectious elimination^36,46,47^. Few CXCR5- and CXCR5+ CD8 T cells produce perforin and CD107a during SLE, but granzyme B is elevated in females at 4 months. Upregulation of granzyme B could be associated with inflammatory tissue destruction. Decreases in perforin and upregulated granzyme B has been associated with multiple cell types in SLE patients and here we further identified these populations in CXCR5+ CD8 T cells^48–52^. CD107a degranulation signaling in these cells may still be functional for delivering perforin to lytic granules but may be unable to overcome SLE’s autoimmune environment in lymph node and spleen.

Our understanding of T cell exhaustion derives from studies of tumor-infiltrating lymphocytes and LCMV. Exhaustion in the context of autoimmunity is less understood, though immunotherapies are being considered for use in treating autoimmune disease^2,53,54^. We unveil a new view to the exhaustion topic in SLE highlighting shifts in CXCR5- and CXCR5+ T cell inhibitory marker expression (PD-1, Tim3, CTLA4, and Lag3). Interestingly, these two CD8 and CD4 T cell populations express different combinations and levels of inhibitory molecules. PD-1 and Tim3 are highly expressed singularly on CXCR5- and CXCR5+ CD8 and CD4 T cells indicating early inhibition through PD-1 with potential for terminal exhaustion through the overlapping expression of Tim3 and Lag3 within both peripheral organs. Tim3 expression has been connected to progressive disease kinetics within multiple human and murine autoimmune diseases^44,55,56^. Tim3 is a gatekeeper of homeostasis and tolerance mechanisms whose expression generates inhibitory functionality towards IFNγ-producing CD8 T cells like the CD8 T cells in this study^45,57^. In autoimmune disease, Tim3 and Lag3 co-expression may provide inhibitory signals that promote tolerance. Lag3 expression restrains autoreactive T cell responsiveness in non-obese diabetic mice^2^. From these results we propose that CXCR5- and CXCR5+ T cell populations are highly functional promoters of autoimmune disease with a small proportion becoming exhausted due to their recognition of self-reactivity. Previous studies have shown that administration of the Tim3 ligand, galectin-9, reduced IFNγ T_H_1 cell numbers^57^. The upregulated expression of Tim3 on CXCR5- and CXCR5+ T cells in this study may be the result of inducing apoptosis by terminal exhaustion and end-stage disease.

CTLA4 expression is an indicator of early exhaustion but has an even greater significance in autoimmune disease. CTLA-4-deficient mice develop severe autoimmune lymphoproliferation indicating regulation of T cell autoimmune progression by CTLA4^58^. CXCR5- and CXCR5+ CD4 T cells in SLE revealed more frequency expression of CTLA4 and Lag3 compared to CD8 T cell populations. Increased B cell production of autoantibodies and inflammatory disease progression has been attributed to CD4 T cells in SLE^34^. CTLA4 depletion is also associated with the upregulation of IFNγ in autoimmune disease^59^. Higher expression of CTLA4 is indicative of lowered T cell activation in CD4 T cells of both peripheral organs suggesting that mechanisms of inhibition are occurring in SLE but unable to overcome the lymphoproliferative environment^60^. The CXCR5- and CXCR5+ CD4 T cells may also contain a population of regulatory T cells that often express CTLA4, though Tregs were not evaluated within this paper. In autoimmune settings Tregs can acquire effecter function due to foxp3 instability^61^.

Contrasting rates of disease between males and females is a common characteristic of autoimmunity. In this study, we identified differences in exhaustion populations but not frequencies associated with sex in MRL/lpr mice. This is profound considering males have a decreased rate of SLE progression in this mouse model and in humans. Little is known about the origins of autoimmune sex bias, but hormones may influence exhaustion upregulation in SLE. In males and females, decreased androgen availability is thought to be protective through upregulating PD-1 on T cells^62^. Though low androgen may also be a result of highly inflammatory autoimmune environments to decrease anti-inflammatory effects of testosterone^63^. In bladder tumor microenvironments, androgen has been shown to upregulate early CD8 T cell dysfunction in T infiltrating lymphocytes of male and female mice leading to increased tumor burden^64^. These studies verify a novel role for androgen signaling in the induction of T cell exhaustion across autoimmune disease and cancer. Overall, the frequency of CXCR5- and CXCR5+ CD8 T cells expressing exhaustion markers was elevated in MRL/lpr mice at all timepoints in males and females, but with different markers, varied timing, and shifts in significance.

## Abbreviations

CTLA4: cytotoxic T-lymphocyte-associated protein 4, CD152
HBV: hepatitis B virus
IFNγ: interferon gamma
IL-1β: interleukin-1 beta
Lag3: lymphocyte activation protein 3, CD223
LCMV: lymphocytic choriomeningitis virus
MRL/MpJ: MRL/MpJ-*Fas^lpr^,* MRL, Murphy Roths Large
MRL/lpr: MRL/MpJ-Fas^lpr^/J, lymphoproliferation
PD-1: programmed cell death protein 1, CD279
SIV: simian immunodeficiency virus
SLE: systemic lupus erythematosus
Tex: terminal exhaustion
Tfh: T follicular helper
Tim3: T-cell immunoglobulin and mucin domain 3, CD366
TNFα: tumor necrosis factor alpha
Tpex: pre-exhausted

## Author Contributions

CNT conceptualization, literature evaluation, experimental investigation, original draft writing, generated and visualized figures. JCSA formal analysis and original draft writing. OAD formal analysis. NH and ARQ experimental investigation. KDCJ original draft review and supervision. KKH conceptualization, writing and review, visualization, funding acquisition, and supervision.

## Acknowledgments

We thank the UC Merced Department of Animal Research Services staff for animal husbandry care and Dr. David Gravano, Science Director of the UC Merced Stem Cell Instrumentation Foundry, for assistance in flow cytometry. Also, Dr. Oscar Davalos for his assistance with bioinformatics and knowledge of T cell exhaustion.

## ORCID iDs

Oscar A. Davalos https://orcid.org/0000-0002-9608-5701

