## Supplemental figures for "T cell exhaustion dynamics in systemic autoimmune disease"

Turner et al. 2023 Supplemental Figure 1

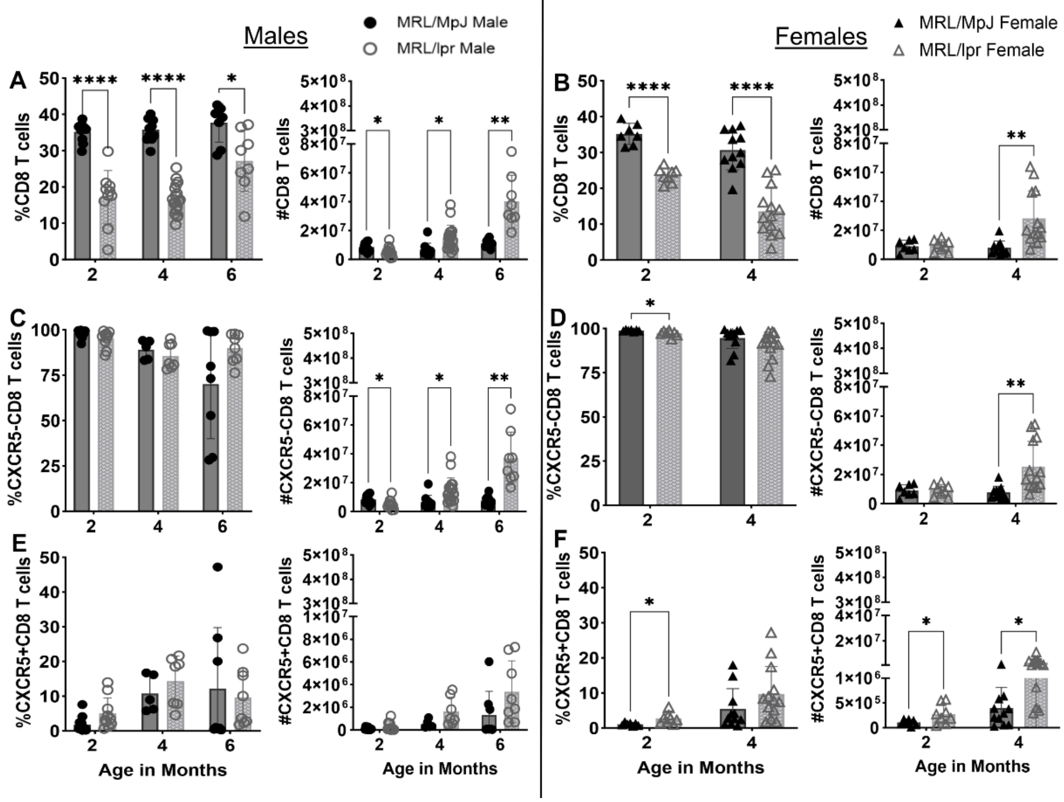


**Supplemental Figure 1. CXCR5- and CXCR5+ CD8 T cell expansion in SLE spleen.** (**A-B**) Frequency and total numbers of all CD8+ T cells across 2, 4, 6 months in male and female MRL/MpJ and MRL/lpr mice. (**C-D**) Frequency and total numbers of CXCR5-CD8 T cells across 2, 4, 6 months in males and female mice. (**E-F**) Frequency and total numbers of CXCR5+CD8 T cells across 2, 4, 6 months in males and female mice. All frequencies and total numbers determined by flow cytometry and cell counting. Each symbol represents one animal and data is representative of 1-5 independent experiments per comparison. Males are denoted by circles and females by triangles. Statistics were performed relative to indicated controls with unpaired Student *t* test. *p < 0.05, **p < 0.01, ****p ≤ 0.0001.

Turner et al. 2023 Supplemental Figure 2

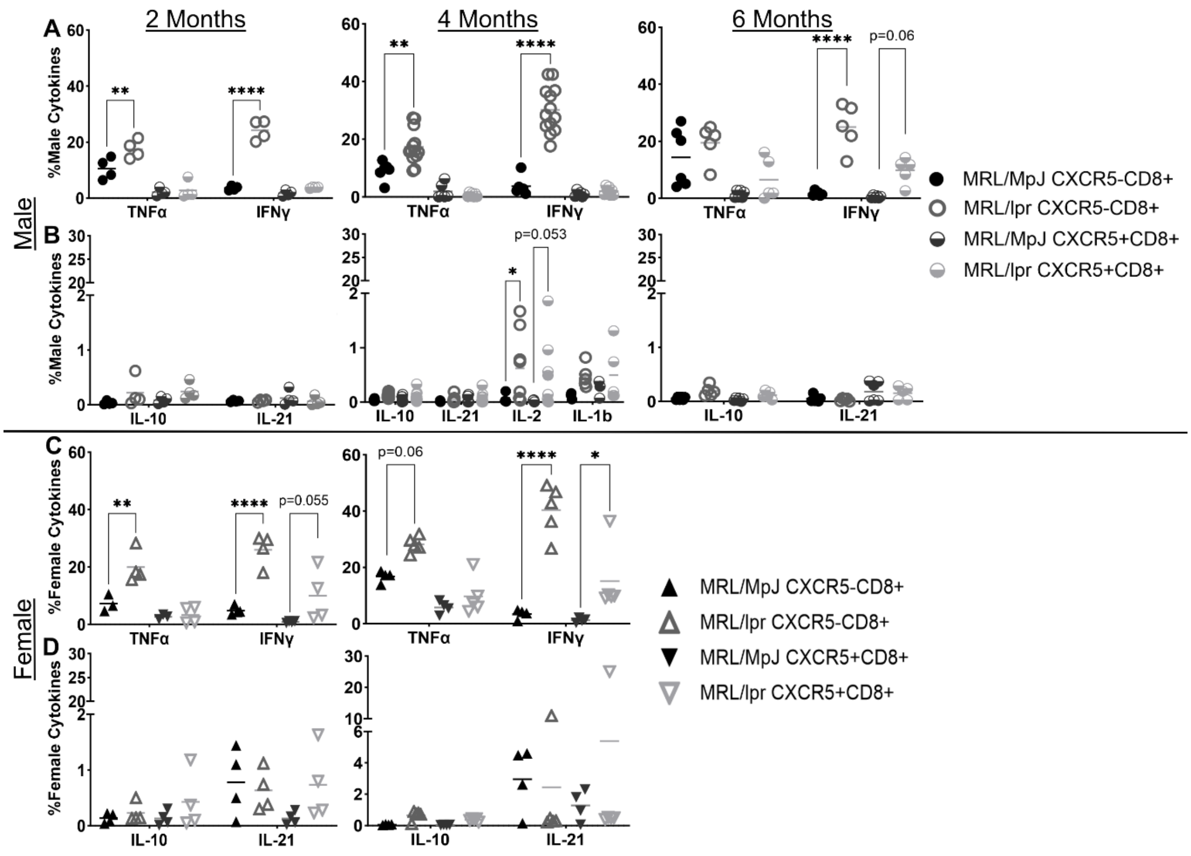


**Supplemental Figure 2. SLE cytokine expression across lifespan on CXCR5- and CXCR5+ CD8 T cells from spleen.** (**A**) Frequency of TNFα and IFNγ producing CXCR5- and CXCR5+ CD8 T cells from male MRL/MpJ and MRL/lpr mice at 2, 4, and 6 months of age. (**B**) Frequency of IL-10 and IL-21 producing CXCR5- and CXCR5+ CD8 T cells from male mice at 2-6 months of age. Frequency of IL-2 and IL-1β producing CXCR5- and CXCR5+ CD8 T cells from male mice at 4 months of age. (**C**) Frequency of TNFα and IFNγ producing CXCR5- and CXCR5+ CD8 T cells from female MRL/MpJ and MRL/lpr mice at 2, 4, and 6 months of age. (**D**) Frequency of IL-10 and IL-21 producing CXCR5- and CXCR5+ CD8 T cells from female mice at 4 months of age. All frequencies determined by flow cytometry. Each symbol represents a different animal and data is representative of 2-5 independent experiments per comparison. Males are denoted by circles and females by triangles. Statistics: ANOVA relative to indicated controls with multiple comparisons and Tukey correction. $^*p < 0.05$, $^{**}p < 0.01$, $^{****}p \le 0.0001$.

**Turner et al. 2023 Supplemental Figure 3**

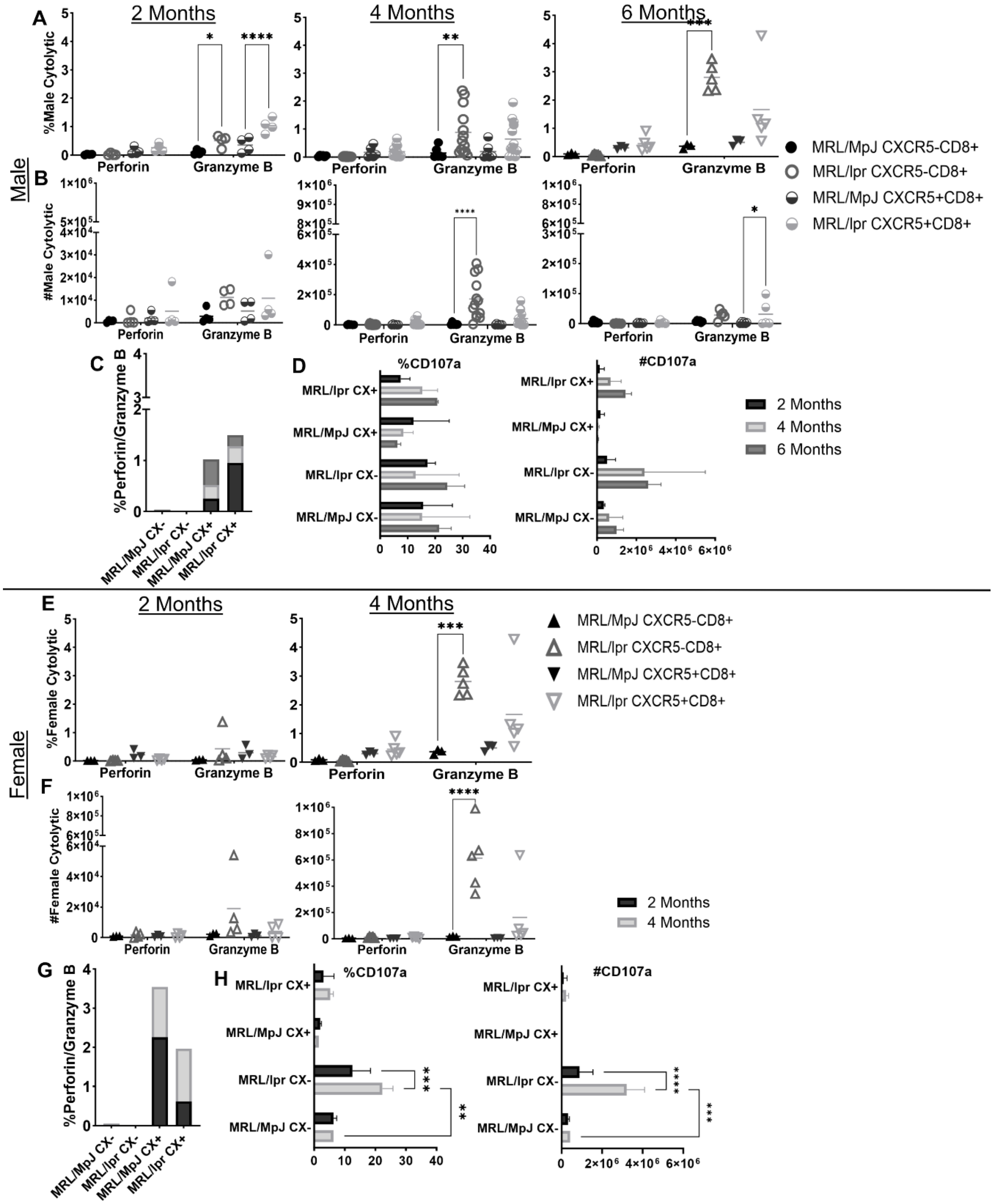


**Supplemental Figure 3. CXCR5- and CXCR5+ CD8 T cell effector molecule production in SLE spleen.** (**A-B**) Frequency and total numbers of perforin and granzyme B producing CXCR5- and CXCR5+ CD8 T cells from male MRL/MpJ and MRL/lpr mice at 2, 4, and 6 months of age. (**C**) Amount of perforin to granzyme B production in male MRL/MpJ versus MRL/lpr CXCR5- and CXCR5+ CD8 T cells. (**D**) Frequency and total number of CD107a degranulation in male

MRL/MpJ and MRL/lpr CXCR5- and CXCR5+ CD8 T cells. (**E-F**) Frequency and total numbers of perforin and granzyme B producing CXCR5- and CXCR5+ CD8 T cells from female MRL/MpJ and MRL/lpr mice at 2, 4, and 6 months of age. (**G**) Amount of perforin to granzyme B production in female MRL/MpJ versus MRL/lpr CXCR5- and CXCR5+ CD8 T cells. (**H**) Frequency and total number of CD107a degranulation in female MRL/MpJ and MRL/lpr CXCR5- and CXCR5+ CD8 T cells. Frequency and total numbers determined by flow cytometry and cell counting. Each symbol represents a different animal and data is representative of 2-5 independent experiments per comparison. Males are denoted by circles and females by triangles. Statistics: ANOVA relative to indicated controls with multiple comparisons and Tukey correction. *p < 0.05, **p < 0.01, ***p ≤ 0.001, ****p ≤ 0.0001.

**Turner et al. 2023 Supplemental Figure 4**

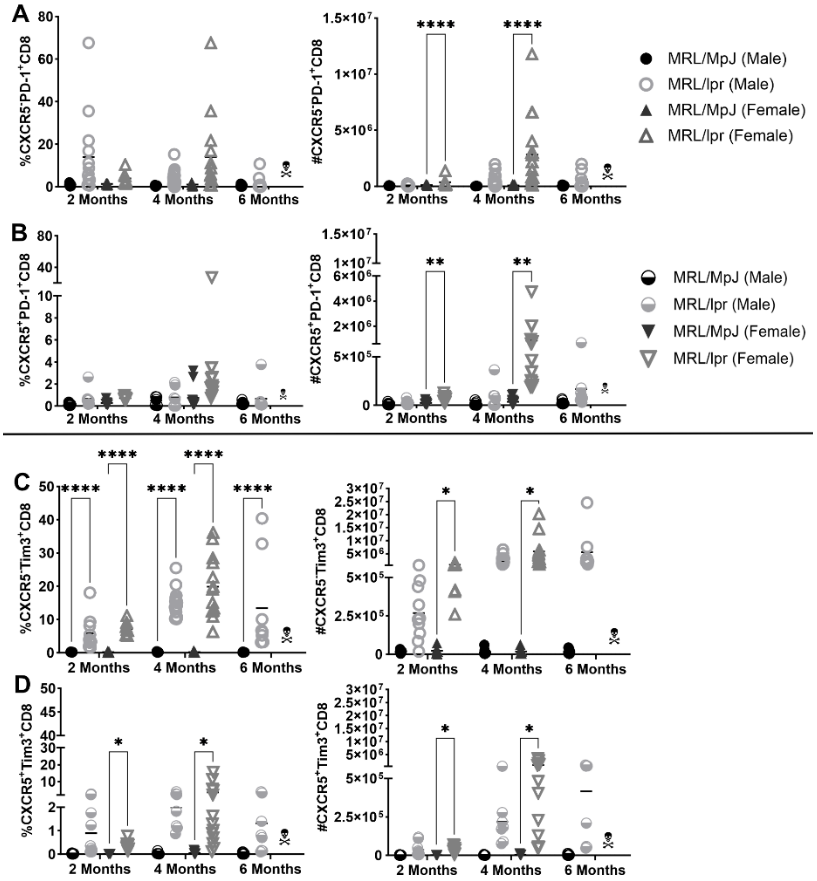


**Supplemental Figure 4. PD-1 and Tim3 expression on CXCR5- and CXCR5+ CD8 T cells from spleen.** CD8 T cells gated on CXCR5 and PD-1 expression on live $B220^-CD11c^-CD11b^-GR\text{-}1^-$ CD8 T cells from 2, 4, and 6-month male and female MRL/MpJ and MRL/lpr mice. (**A**) Frequency of PD-1 expressing CXCR5- CD8 T cells from 2, 4, and 6-month male and female MRL/MpJ and MRL/lpr mice. (**B**) Frequency of PD-1 expressing CXCR5+ CD8 T cells from 2, 4, and 6-month male and female MRL/MpJ and MRL/lpr mice. (C) Frequency of Tim3 expressing CXCR5- CD8 T cells from 2, 4, and 6-month male and female MRL/MpJ and MRL/lpr mice. (**D**) Frequency of Tim3 expressing CXCR5+ CD8 T cells from 2, 4, and 6-month male and female MRL/MpJ and MRL/lpr mice. Each symbol represents a different animal and data is representative of 2-5 independent experiments per comparison. Males are denoted by circles and females by triangles. Statistics: ANOVA relative to indicated controls with multiple comparisons and Tukey correction. *p < 0.05, **p < 0.01, ****p ≤ 0.0001.

**Turner et al. 2023 Supplemental Figure 5**

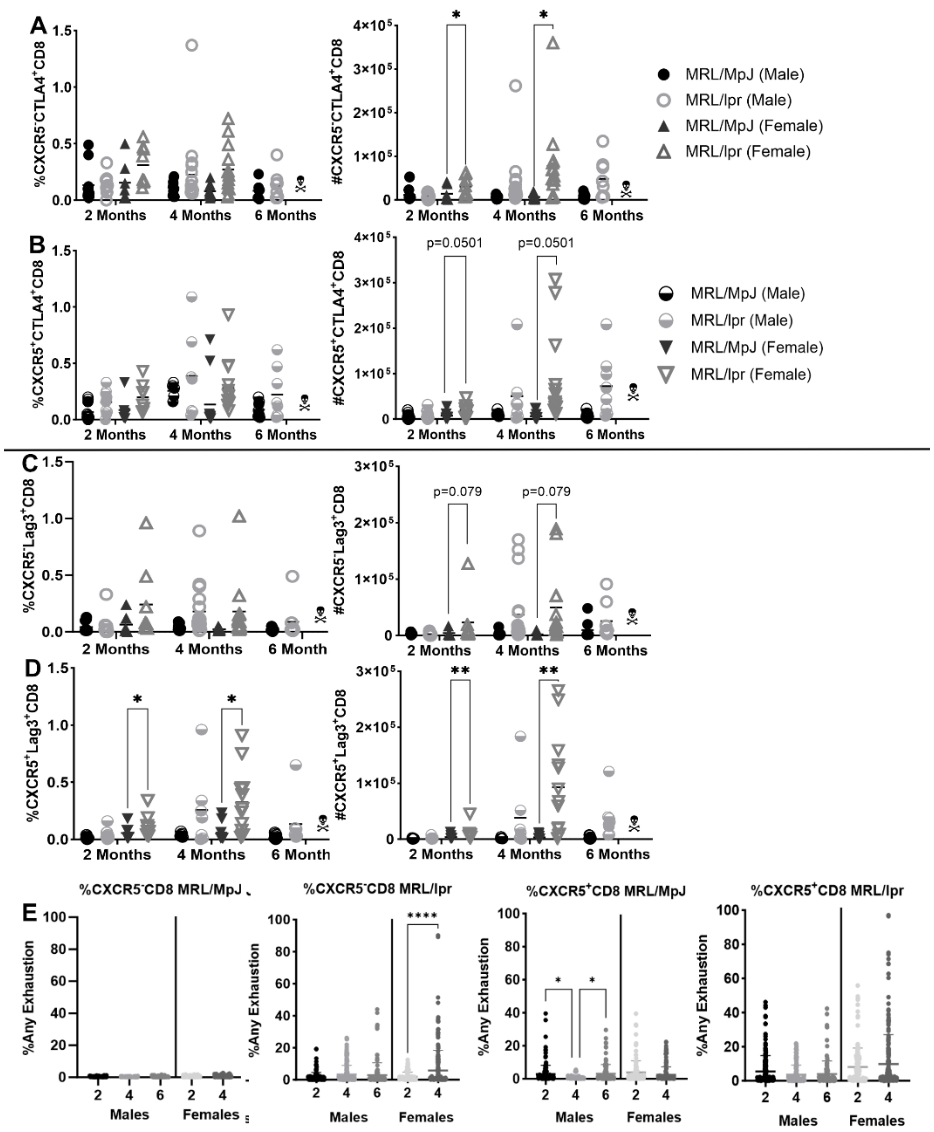


**Supplemental Figure 5. CTLA4 and Lag3 expression on CXCR5- and CXCR5+ CD8 T cells from spleen.** CD8 T cells gated on CXCR5 and CTLA4 expression on live B220$^-$CD11c$^-$CD11b$^-$ GR-1$^-$ CD8 T cells from 2, 4, and 6-month male and female MRL/MpJ and MRL/lpr mice. (**A**) Frequency of CTLA4 expressing CXCR5- CD8 T cells from 2, 4, and 6-month male and female MRL/MpJ and MRL/lpr mice. (**B**) Frequency of CTLA4 expressing CXCR5+ CD8 T cells from 2, 4, and 6-month male and female MRL/MpJ and MRL/lpr mice. (**C**) Frequency of Lag3 expressing CXCR5- CD8 T cells from 2, 4, and 6-month male and female MRL/MpJ and MRL/lpr mice. (**D**) Frequency of Lag3 expressing CXCR5+ CD8 T cells from 2, 4, and 6-month male and female MRL/MpJ and MRL/lpr mice. (**E**) Any individual and double exhaustion marker frequencies were added together for each MRL/MpJ and MRL/lpr male and female mouse. The average frequency was graphed to show all exhaustion marker expression across 2-6 months in male and female mice. Each symbol represents a different animal and data is representative of 2-5 independent experiments per comparison. Males are denoted by circles and females by triangles. Statistics: ANOVA relative to indicated controls with multiple comparisons and Tukey correction. *p < 0.05, **p < 0.01, ****p ≤ 0.0001.

**Turner et al. 2023 Supplemental Figure 6**

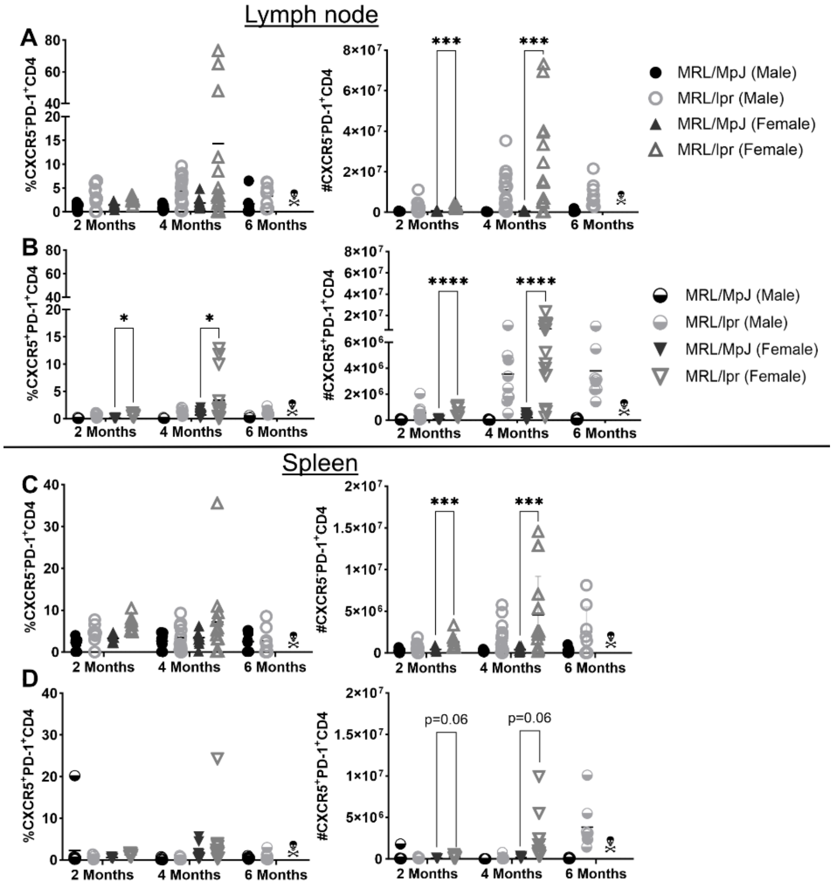


**Supplemental Figure 6. PD-1 expression on CXCR5- and CXCR5+ CD4 T cells from lymph node and spleen.** CD4 T cells gated on CXCR5 and PD-1 expression on live B220$^-$CD11c$^-$CD11b$^-$ GR-1$^-$ CD8 T cells from 2, 4, and 6-month male and female MRL/MpJ and MRL/lpr mice. (**A**) Frequency of PD-1 expressing CXCR5- CD4 T cells from lymph nodes of 2, 4, and 6-month male and female MRL/MpJ and MRL/lpr mice. (**B**) Frequency of PD-1 expressing CXCR5+ CD4 T cells from lymph nodes of 2, 4, and 6-month male and female MRL/MpJ and MRL/lpr mice. (**C**) Frequency of PD-1 expressing CXCR5- CD4 T cells from spleens of 2, 4, and 6-month male and female MRL/MpJ and MRL/lpr mice. (**D**) Frequency of PD-1 expressing CXCR5+ CD4 T cells from spleens of 2, 4, and 6-month male and female MRL/MpJ and MRL/lpr mice. Each symbol represents a different animal and data is representative of 2-5 independent experiments per comparison. Males are denoted by circles and females by triangles. Statistics: ANOVA relative to indicated controls with multiple comparisons and Tukey correction. *p < 0.05, ***p ≤ 0.001, ****p ≤ 0.0001.

**Turner et al. 2023 Supplemental Figure 7**

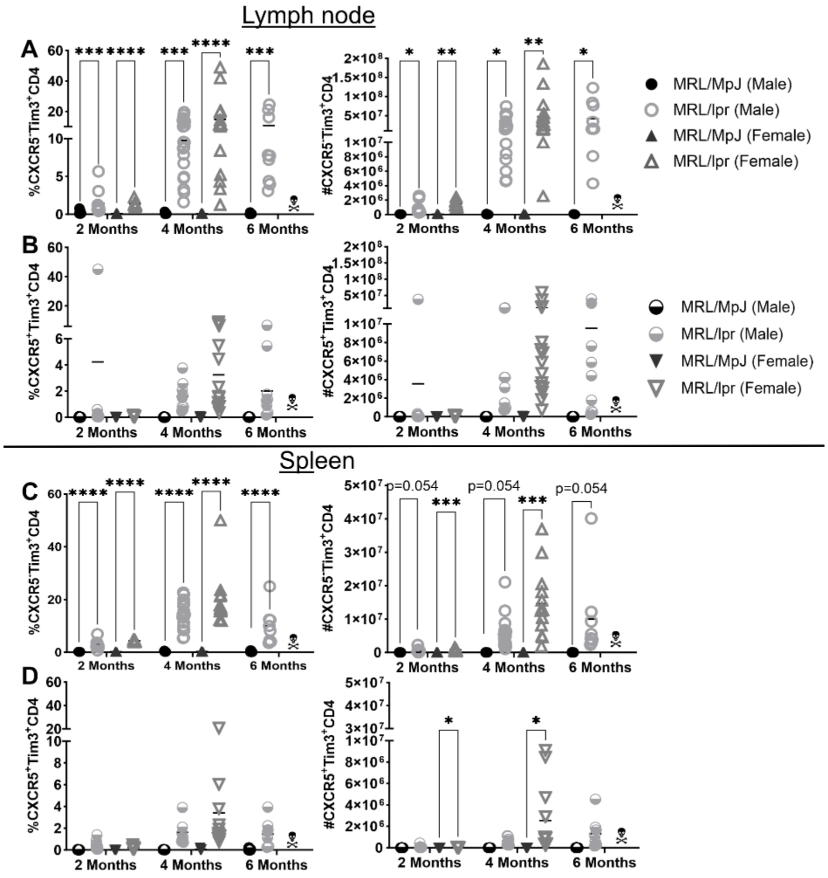


**Supplemental Figure 7. Tim3 expression on CXCR5- and CXCR5+ CD4 T cells from lymph node and spleen.** CD4 T cells gated on CXCR5 and Tim3 expression on live B220$^-$CD11c$^-$CD11b$^-$ GR-1$^-$ CD8 T cells from 2, 4, and 6-month male and female MRL/MpJ and MRL/lpr mice. (**A**) Frequency of Tim3 expressing CXCR5- CD4 T cells from lymph nodes of 2, 4, and 6-month male and female MRL/MpJ and MRL/lpr mice. (**B**) Frequency of Tim3 expressing CXCR5+ CD4 T cells from lymph nodes of 2, 4, and 6-month male and female MRL/MpJ and MRL/lpr mice. (**C**) Frequency of Tim3 expressing CXCR5- CD4 T cells from spleens of 2, 4, and 6-month male and female MRL/MpJ and MRL/lpr mice. (**D**) Frequency of Tim3 expressing CXCR5+ CD4 T cells from spleens of 2, 4, and 6-month male and female MRL/MpJ and MRL/lpr mice. Each symbol represents a different animal and data is representative of 2-5 independent experiments per comparison. Males are denoted by circles and females by triangles. Statistics: ANOVA relative to indicated controls with multiple comparisons and Tukey correction. *$p < 0.05$, **$p < 0.01$, ***$p \leq 0.001$, ****$p \leq 0.0001$.

**Turner et al. 2023 Supplemental Figure 8**

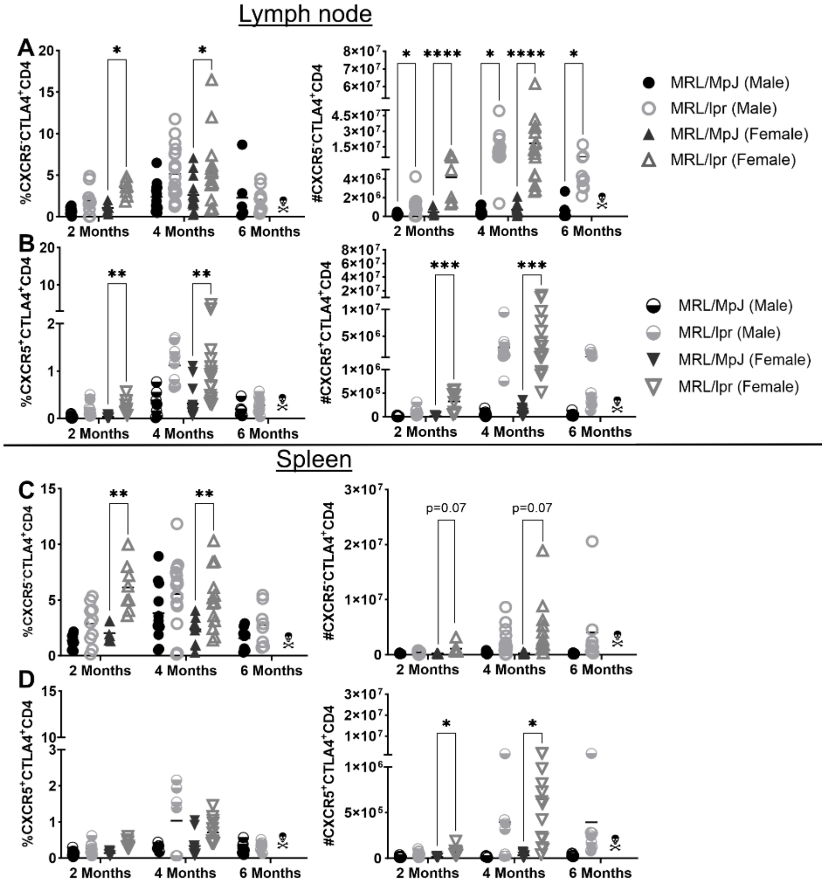


**Supplemental Figure 8. CTLA4 expression on CXCR5- and CXCR5+ CD4 T cells from lymph node and spleen.** CD4 T cells gated on CXCR5 and CTLA4 expression on live B220$^-$CD11c$^-$ CD11b$^-$GR-1$^-$ CD8 T cells from 2, 4, and 6-month male and female MRL/MpJ and MRL/lpr mice. (**A**) Frequency of CTLA4 expressing CXCR5- CD4 T cells from lymph nodes of 2, 4, and 6-month male and female MRL/MpJ and MRL/lpr mice. (**B**) Frequency of CTLA4 expressing CXCR5+ CD4 T cells from lymph nodes of 2, 4, and 6-month male and female MRL/MpJ and MRL/lpr mice. (**C**) Frequency of CTLA4 expressing CXCR5- CD4 T cells from spleens of 2, 4, and 6-month male and female MRL/MpJ and MRL/lpr mice. (**D**) Frequency of CTLA4 expressing CXCR5+ CD4 T cells from spleens of 2, 4, and 6-month male and female MRL/MpJ and MRL/lpr mice. Each symbol represents a different animal and data is representative of 2-5 independent experiments per comparison. Males are denoted by circles and females by triangles. Statistics: ANOVA relative to indicated controls with multiple comparisons and Tukey correction. *$p < 0.05$, **$p < 0.01$, ***$p \leq 0.001$, ****$p \leq 0.0001$.

**Turner et al. 2023 Supplemental Figure 9**

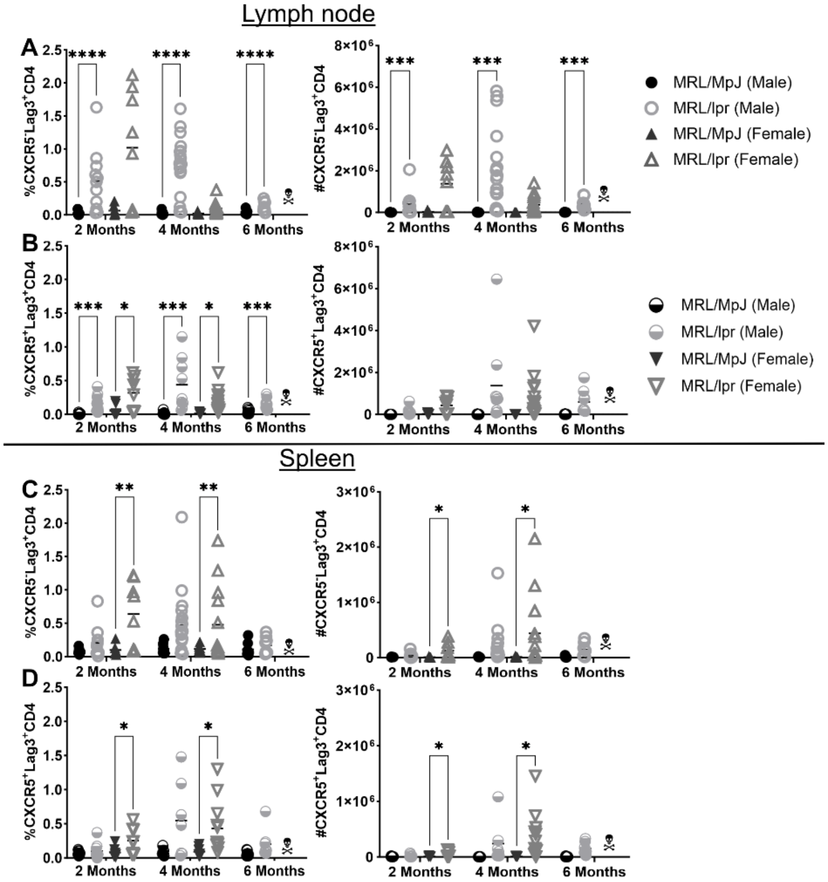


**Supplemental Figure 9. Lag3 expression on CXCR5- and CXCR5+ CD4 T cells from lymph node and spleen.** CD4 T cells gated on CXCR5 and Lag3 expression on live B220⁻CD11c⁻CD11b⁻ GR-1⁻ CD8 T cells from 2, 4, and 6-month male and female MRL/MpJ and MRL/lpr mice. (**A**) Frequency of Lag3 expressing CXCR5- CD4 T cells from lymph nodes of 2, 4, and 6-month male and female MRL/MpJ and MRL/lpr mice. (**B**) Frequency of Lag3 expressing CXCR5+ CD4 T cells from lymph nodes of 2, 4, and 6-month male and female MRL/MpJ and MRL/lpr mice. (**C**) Frequency of Lag3 expressing CXCR5- CD4 T cells from spleens of 2, 4, and 6-month male and female MRL/MpJ and MRL/lpr mice. (**D**) Frequency of Lag3 expressing CXCR5+ CD4 T cells from spleens of 2, 4, and 6-month male and female MRL/MpJ and MRL/lpr mice. Each symbol represents a different animal and data is representative of 2-5 independent experiments per comparison. Males are denoted by circles and females by triangles. Statistics: ANOVA relative to indicated controls with multiple comparisons and Tukey correction. *p < 0.05, **p < 0.01, ***p ≤ 0.001, ****p ≤ 0.0001.

**Turner et al. 2023 Supplemental Figure 10**

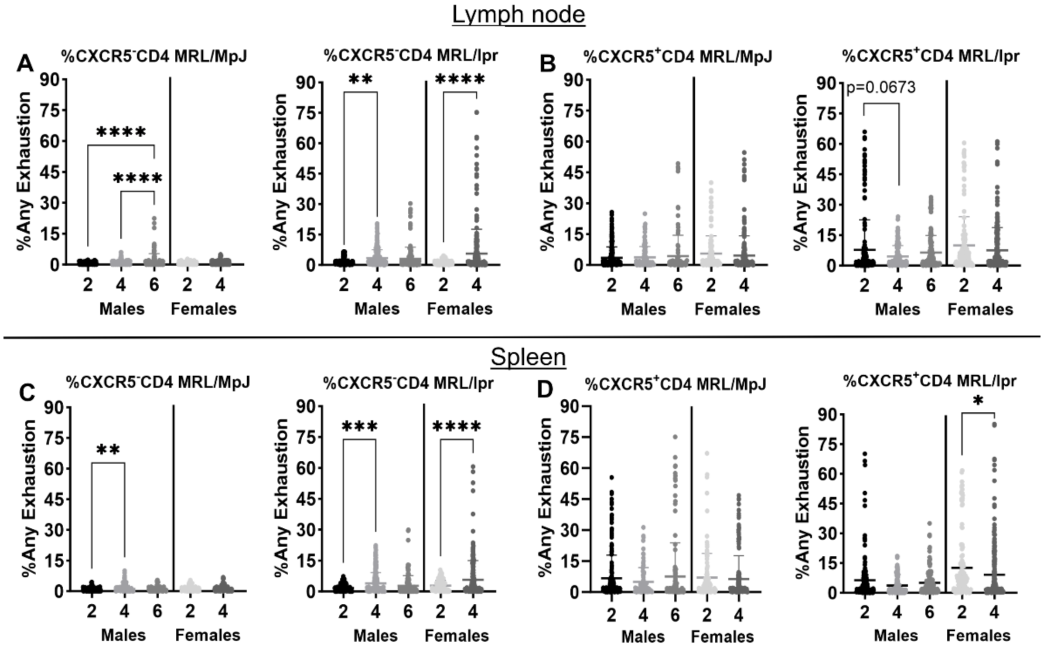


**Supplemental Figure 10. CXCR5- and CXCR5+ CD4 T cells upregulate exhaustion markers in SLE.** CD4 T cells gated on live $B220^{-}CD11c^{-}CD11b^{-}GR\text{-}1^{-}$ CD8 T cells from 2, 4, and 6-month male and female MRL/MpJ and MRL/lpr mice. (**A-D**) Any individual and double exhaustion marker frequencies were added together for each MRL/MpJ and MRL/lpr male and female mouse. The average frequency was graphed to show all exhaustion marker expression across 2-6 months in male and female lymph nodes and spleens. Each symbol represents a different animal and data is representative of 2-5 independent experiments per comparison. Males are denoted by circles and females by triangles. Statistics: ANOVA relative to indicated controls with multiple comparisons and Tukey correction. *p < 0.05, **p < 0.01, ***p ≤ 0.001, ****p ≤ 0.0001.
